# Solo songs, duets, and territory defence across seasons in female Galápagos Yellow Warblers (*Setophaga petechia aureola*)

**DOI:** 10.1101/2025.06.02.657377

**Authors:** Alper Yelimlieş, Katherine Albán Morales, Çağlar Akçay, Sonia Kleindorfer

**Affiliations:** Konrad Lorenz Research Center for Behavior and Cognition, Grünau im Almtal, 4645 Austria; Department of Behavioral and Cognitive Biology, University of Vienna, Vienna, 1030 Austria; School of Life Sciences, Anglia Ruskin University, Cambridge, UK

**Keywords:** Female song, Aggression, *Setophaga petechia aureola*, Bird song function

## Abstract

While the function of bird song has been well studied in male songbirds, the function of female song is less well understood. This is partly due to a historical view of females occupying a passive role compared with males, which led to ignoring female song even in some well-studied species. We report one such case in yellow warblers (*Setophaga petechia*), as no other study investigated female song in 35 years after the first documentation of solo female songs. We interrogate the seasonal patterns and functions of female song in Galápagos yellow warblers (*Setophaga petechia aureola*), in which females perform solo song and produce duets with their male social partners. We carried out simulated territory intrusions by broadcasting male, female, or duet songs during breeding and non-breeding seasons, and conducted a territory retention survey for over a year. We measured the association between aggressive response and singing behaviour, sex-specific patterns of response, and territory retention across years. Females sang mostly during the non-breeding season and predominantly in male-led duets. Although females were strongly aggressive towards female song playback, they gave the weakest singing response towards them. There was no association between female aggressive behaviour and song output in response to a simulated intruder. Moreover, the probability of territory retention across years was not explained by song output or aggression in response to intruders, though evidence for this was weak due to the small sample size. We suggest that female song in this year-round resident island system does not function for territory defense or intrasexual competition, but may have other functions.

**Highlights:**

- Galápagos Yellow Warbler females produce solo songs and duets with their male partners.
- Female song and aggression were mainly restricted to the non-breeding season.
- While females were equally aggressive to all intruders, they had the lowest song rate in response to females.
- In contrast to males, aggressive behaviour didn’t predict song output in females.
- Female singing behaviour did not increase the probability of territory retention.

---

The study of sexual selection has largely focused on male ornaments while neglecting females. This biased focus has been compounded historically by a longstanding view of females occupying a passive role compared with males (Ah-King, 2022b; Dougherty, 2021; Hrdy, 1986), which consequently has impeded the progress in our understanding of the evolutionary history and significance of female behavior. More recently, this view is being replaced by females as active participants of inter- and intrasexual selection with the inclusion of feminist perspectives into research (Ah-King, 2022a; Tang-Martínez, 2020).

Research on bird song is a good example of this historical trajectory. Although females were long known to sing in some species, female song was viewed as an exceptional phenomenon, mostly occurring in the tropics or as a byproduct of abnormal testosterone levels (Catchpole & Slater, 2008). Increased attempts to address this bias in the last decades revealed that singing females are not exceptions; rather, female song is estimated to be an ancestral trait in songbirds (Odom et al., 2014) and present in 59% of the songbird species (Odom et al., 2024). Despite the significant progress, the current state of knowledge is nonetheless incomplete, as we lack sex specific singing information on 73% of the songbird species (Odom & Benedict, 2018).

Increased research on female song has revealed both convergent and divergent patterns of singing and song structure between sexes (Evans & Kleindorfer, 2016; Liu et al., 2024; Magoolagan et al., 2019; Moyer et al., 2025; Patchett et al., 2021; Sierro et al., 2022). The divergent structural patterns suggest functions of singing may be sex specific in at least some species (Morton, 1977); therefore, a complete understanding of bird song function requires separate investigation of both males and females (Barros et al., 2024).

The ecological significance of song is primarily explained by its dual function: mate attraction and territory defence (Catchpole & Slater, 2008). To date, these functions have mostly been tested in males, who sing to attract mates and defend their resources against same sex intruders (Eriksson & Wallin, 1986; Krebs et al., 1978). Studies on female song generally supported the same dual function (Beletsky, 1983; Krieg & Getty, 2016; Langmore et al., 1996). Additionally, females of some species were shown to use song in within-pair (Beletsky, 1983; Halkin, 1997; Rose et al., 2019) or parent-offspring communication (Ritchison, 1983). Barros et al. (2024) integrated the relative support for these five main functions of female solo singing, reviewing 147 studies. They found that the most supported function was territory defense, followed by intrapair communication and intrasexual competition. In line with this finding, territoriality was also shown to predict the presence of female singing in multi-species analyses (Mikula et al., 2020; Odom et al., 2024). Moreover, both studies found that among the species with female song, duetting (coordinated singing within a pair) was more common in year-round territorial species than seasonally territorial species.

The yellow warbler (*Setophaga petechia*) is a suitable species for testing the territoriality-related functions of female solo song. Hobson and Sealy (1990) previously reported that female song is rare and is only produced as solo song in the migratory North American subspecies. Their observations suggested that territorial resource defense against conspecific females during the early breeding season could be the main function of female song. Although the yellow warbler is a well-studied species, including in the context of male song and other female vocalizations (Beebee, 2002, 2004; Ficken & Ficken, 1965; Gill & Sealy, 2004; Spector, 1991; Spector et al., 1989; Studd & Robertson, 1985; Yezerinac & Weatherhead, 1997), there have been no follow-up studies on female song, likely because it is rarely observed in the migratory populations. The current study started with our observation that, in contrast to the migratory populations in the non-migratory Galápagos subspecies of yellow warblers (*Setophaga petechia aureola*), females commonly produce solo songs as well as duets with males.

We test two non-mutually exclusive hypotheses for female song in the Galápagos yellow warbler, namely, intrasexual competition and territory defense. To this aim, we carried out simulated territory intrusion experiments with focal pairs in breeding and non-breeding seasons by broadcasting solo male, solo female, and duet songs. We also conducted separate surveys to compare the probability of territory retention depending on the song rate from June 2023 to March 2025. If female song functions in intrasexual competition, we would expect resident females to respond with more aggression and singing to intruder females than males. If female song functions in territory defense (defense of physical resources against both males and females) we predict that 1) playback of female song will elicit equal aggressive responses from both the male and the female and responses will be comparable to playback of male song; 2) resident female aggressive responses in response to playback of either male or female song will be associated with her song output. Additionally, we predict that 3) pairs will respond more strongly to the duetting intruders as an intruding pair is assumed to be a stronger threat to territory ownership than a single bird of either sex. Finally, we predict that females that are singing more and are more aggressive will be more likely to retain their territory across years. We also report on the occurrence of female singing during the breeding and non-breeding seasons.

## Methods

### Study species

We studied the Galápagos yellow warblers on Floreana island in the Galápagos archipelago. Phylogenetic estimates show that the *aureola* subspecies of the yellow warbler split from the mainland and colonized the Galápagos Archipelago approximately 300.000 years ago (Chaves et al., 2012). Unlike the commonly studied North American subspecies, the Galápagos subspecies is sedentary and territorial year-round (Snow, 1966). Males and females form long-term pair bonds, and their territories are stable across years (see results). Breeding occurs typically after the onset of the rains in January and extends throughout the rainy season, which can be variable on the Galápagos but typically extends until April. Snow (1966) observed them breeding from December to April on Santa Cruz Island. Pairs can raise one or more broods per season, with incubation and feeding each lasting about ten to fourteen days, followed by post-fledging feeding of young. Females build the nest and incubate the eggs; both parents feed the chicks.

In the field, sexes can be distinguished by their song and plumage with ease. Males have rufous crowns and rufous chest streakings over their yellow and olivaceous plumage. In contrast, females lack the crown and have little to no streakings (see Figure 1 for illustrations). Female songs usually have more repetitive elements and sound harsher than male songs to the human ear. Moreover, males and females often combine their songs into polyphonous duets (see Figure 1 for spectrogram examples).

**Figure 1.**
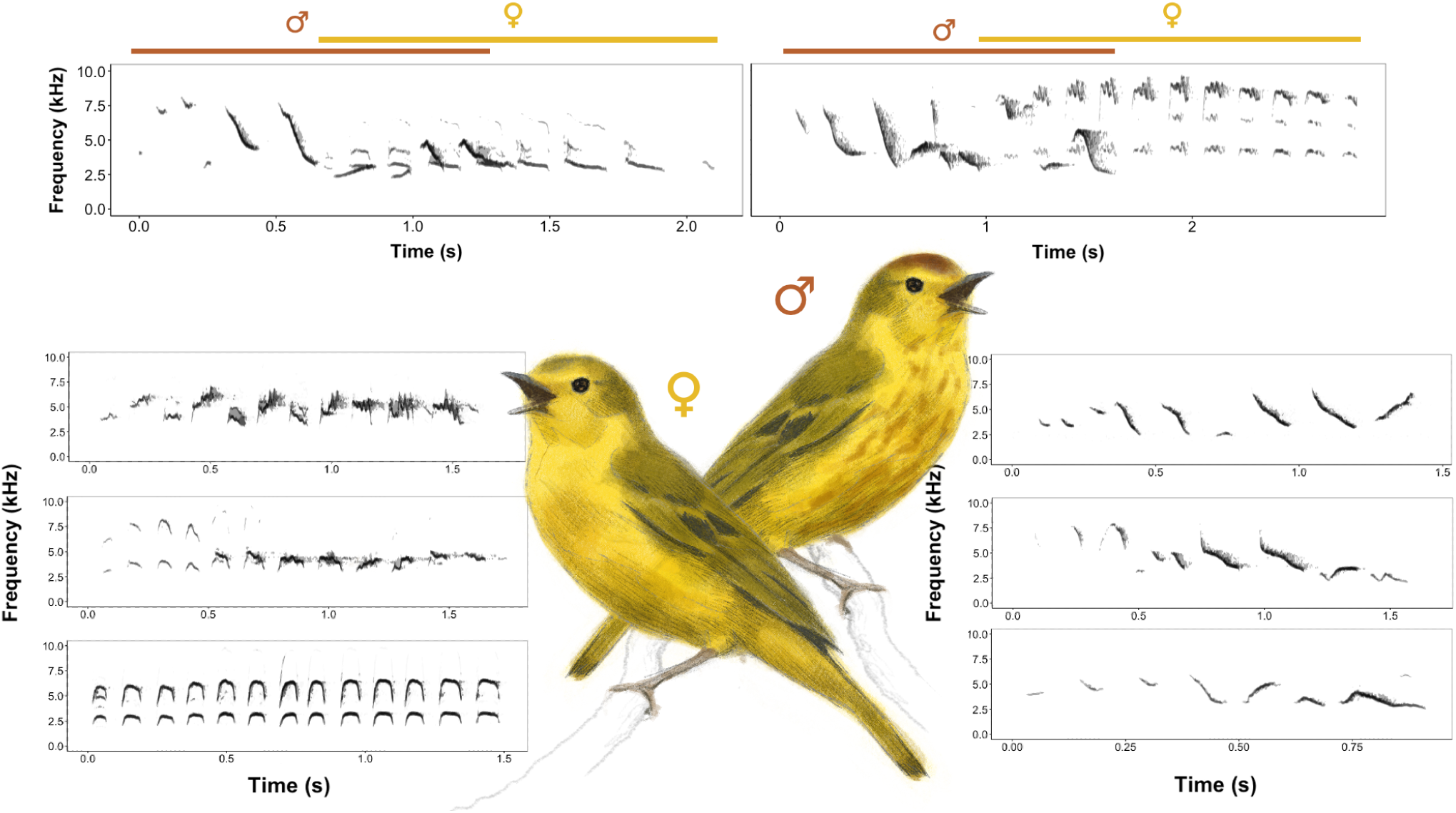
Illustration of female and male yellow warblers and their respective songs. On top, two duets are shown in spectrograms with male parts marked in red and female parts marked in yellow. On the left are three female songs and on the right are three different male songs. All displayed songs are obtained from different individuals. Spectrograms are created using 512 samples per window and 99% overlap. Illustrations are courtesy of Lena Gies.

### Study site

We studied yellow warblers between June and July 2023 (non-breeding season) and between January and March 2024 (breeding season) on multiple sites on Floreana Island including Cerro Pajas (−1.295927, −90.456195), Asilo de La Paz (−1.313046, −90.454748), and Puerto Velazco Ibarra (the only settlement on the island, −1.275715, −90.486968) that have been a part of long-term research of our group on songbirds since 2004 (Kleindorfer et al., 2021, see figure S1 for the playback locations).

### Playback experiment

#### Stimulus preparation

To make the playback stimuli, we recorded male and female songs in June 2023 at a 48 kHz sampling rate and 24-bit resolution using a Sennheiser MKE 600 directional microphone attached to a Zoom H5 recorder. From our recordings, we selected 10 high-quality male and female songs from different individuals and further processed them in Audacity version 3.3.3. We first high-pass filtered the recordings above 1500 Hz and then normalized the peak amplitude. Then, we created solo male and solo female stimulus files by repeating these songs at a rate of 6 songs per minute. This corresponds to how males sing during agonistic interactions (Beebee, 2004). For the duet stimuli, we combined the songs of both sexes, whereby the female song followed the male song 0.5 seconds after it had ended. This response delay is within the natural limit of the species’ duets, and we chose to keep it constant across iterations for the sake of simplicity.

#### Experimental procedure

We determined active territories by observing singing posts and the territorial behavior of the pair (described in Hohl et al., 2025). Upon arrival in the territory of a pair, we placed a Sony SRS-XB100 Bluetooth speaker in a central location at 1.5 m above the ground. During the breeding season, we placed the speaker 10 m away from the nest and within the territory to minimize the disturbance to the breeding activity and to make our response variables relative to the speaker’s position more valid. Then, we marked 1 and 3-m distances from the speaker with flagging tape in two directions to assist with our distance estimates. To start the trials, we waited until both sexes were within 20 m of the speaker, confirmed either by sight or hearing. Then we started the playback period, which consisted of one minute of song playback followed by one minute of silence and another minute of song for a total of three minutes. The broadcast amplitude of the stimuli for both sexes was set to ∼80 dB measured using a sound level meter (Svan 971, Svantek UK LTD, A-weighting, fast setting) at 1 m. During the playback period, we narrated the behavior of the pair and recorded their vocalizations using a Marantz PMD660 recorder with a Sennheiser ME66/K6 shotgun microphone or a Zoom F3 recorder with a Sennheiser MKE 600 shotgun microphone. After the playback period was over, we recorded the vocalizations of the focal pair for another 5 minutes. All observations were done by two experienced field researchers (either AY, CA, or KAM), one was narrating the behavior for the male and the other one narrating the behavior for the female, to ensure all behavior was recorded with as little error as possible.

We tested 17 pairs in the non-breeding season (June-July 2023) and 20 pairs in the breeding season (January-March 2024). In the non-breeding season, both members of eight pairs and one member of three pairs (one male and two females) were color-banded before the experiments; the rest of the birds were not banded. Of the pairs tested in the breeding season, both members of six pairs and a single member of two pairs were banded (one male and one female). In total, eight territories were tested in both seasons. Out of those eight territories, two were occupied by unbanded individuals, the rest were occupied by the same banded pair in both seasons. For our analyses, we assumed the remaining two territories were also occupied by the same individuals in both seasons. This assumption did not affect any of our results qualitatively. For each trial, we used a non-neighboring individuals’ songs and avoided testing neighboring pairs on the same day. Each pair was tested three times with solo male, solo female, and duet song treatments on the same day. The order of trials was randomly chosen (see Supplementary Table 1 for the distribution of treatment orders), and consecutive trials were at least 45 minutes apart for the same pair. All playback trials were done between 6 AM and 11:30 AM.

#### Quantifying aggressive and song response

We used Audacity version 3.3.3 to annotate our recordings of trials. From the annotations, we extracted the number of flights, the closest approach to the speaker, time spent within 5m of the speaker, and latency to respond to playback for the three-minute playback period as proxies for physical aggression. All these variables are commonly used in simulated intrusion experiments of songbirds as indicators of territorial aggression (Searcy et al., 2006). We also annotated the number of songs for each bird and duets for each pair for the 3-minute playback period and the following 5-minute post-playback period. We defined duets as consecutive songs from both sexes in which the response song starts after the initial song, and at the latest two seconds after the initial song ends. If the initiator of the duet sang again, overlapping the responder or within 2 seconds after the end of the response song, we counted these consecutive songs as one duet. For our analyses of song, we first categorized individuals as singers and non-singers based on whether they sang any solo or duet songs in response to any of the treatments in each season.

### Territorial retention surveys

To quantify the relationship between song rate, aggression, and territory retention, we conducted territory surveys. In 2024 and 2025, we visited each territory with banded birds and coded the female as absent or present. More specifically, 18 banded females were observed from 2023 to 2025, and 8 females were observed from 2024 to 2025. In the cases in which the bird was absent, we confirmed that another unbanded individual of the same sex was defending the territory by doing playbacks in various locations around the territory. For our analyses on aggression and song rate, we used 9 of the banded females tested in our playback experiment in the non-breeding season, which were also included in our surveys.

### Data analysis

All statistical analyses were conducted in R version 4.3.3 (R Core Team, 2024). As the aggression variables (number of flights, closest approach distance, and latency to respond, time spent within 5m of the speaker) were correlated with each other, we performed a Principal Component Analysis (PCA) to reduce dimensions using the *prcomp* with centered and scaled variables. We used the *PCAtest* package with 1000 permutations and bootstrap replication to evaluate the significance of the PCA and the resulting PCs (Camargo, 2022). PC1 was significant, accounted for 67.5% (95% CI: 64.2-71.1) of the total variation in the data, and had an eigenvalue of 2.70. All variables had significant loadings on PC1 (factor loadings can be seen in Supplementary Table 2). We took the inverse scores as our aggression variable so that the higher values for this variable indicated higher aggression.

We analyzed the resulting aggression scores (PC1) using a linear mixed model with Gaussian error distribution using the R package *lme4* (Bates et al., 2015). The model included season (breeding, non-breeding), sex (male, female), treatment (male, female, duet), all of their two-way interactions, the three-way interaction between them as the fixed effects, and bird ID nested within pair ID as the random effects. In the initial model, we controlled for the effect of treatment order by adding it as a fixed factor, but as it did not reveal a significant effect, we removed it from the final model.

We first wanted to quantify individuals who sang at least one song in response to any of the treatments in both seasons. To test if the number of singing individuals depends on the season, we performed a chi-square test. We performed the test only for the females because all the males were singing in both seasons.

As our test revealed that females were mostly singing in the non-breeding season (see the results section), we restricted our analyses of the number of solo songs and duets produced during playback and post-playback periods to the non-breeding season. We first calculated descriptive statistics such as the proportion of solo songs each sex produced and the proportion of duets led by either sex. Then, we modeled each of these two response variables using GLMMs with Poisson error distribution and log link function using the package *lme4* (Bates et al., 2015). The number of solo songs model initially included sex, treatment, their interaction, and the treatment order as the fixed effects, and bird ID nested within pair ID as the random effects. As the treatment order effect was not significant, we removed the term from the final model. Because duetting was a pair-level response, the corresponding model included treatment and treatment order as the fixed effects and pair ID as the random effect.

We then tested whether singing is associated with aggression by modeling whether the aggression scores during playback trials predicted songs produced post-trial using a zero-inflated GLMM with Poisson error distribution and log link function using the package *glmmTMB* (Brooks et al., 2017). As females were mainly singing in the non-breeding season, we excluded the breeding season from the model. For the males, both seasons were used. The model had the songs produced after the trial as the response variable, aggression score (PC1), sex, and their interaction as the fixed factors, and bird ID as the random factor. Zero-inflated part of the model included sex as the fixed factor and bird ID as the random factor because we expected zeroes mainly resulting from females.

Finally, we modeled the relationship between song rate and territory retention using the data from our territory surveys. For this, we used a GLM with a binomial error distribution, which had the territory retention in 2025 (0 = lost, 1 = retained) as the response variable average number of songs across the three playback trials as the predictor variable. We also modeled the probability of territory retention as a function of average aggression score over three trials.

We checked model diagnostics for all the models above using the package *DHARMa* (Hartig, 2017). We report predictor estimates, 95% confidence intervals, as well as *χ2* and p-values for overall predictors from type 3 ANOVA using the package *car* (Fox & Weisberg, 2019). When needed, we also report Tukey p-value adjusted post hoc comparisons using the package *emmeans* (Lenth, 2023). We used the package *ggeffects* to create the plot for the relationship between aggression and singing (Lüdecke, 2018).

### Ethical note

This study complies with all current Austrian laws and regulations and was supported by Animal Experiment License Number 66.006/0026-WF/V/3b/2014 issued by the Austrian Federal Ministry for Science and Research (EU Standard, equivalent to the Animal Ethics Board). The subjects were not captured during the experiment, and each trial only took eight minutes, which included two minutes of active playback. The birds returned to their normal activities within a few minutes.

## Results

### Responses to the playback experiment

#### Aggressive behavior

Aggressive responses to simulated intruders differed between treatments, sexes and seasons. There was a significant interaction effect between sex and season on aggression scores (*χ2* = 9.53, *P* < 0.001), with less aggression in females in the breeding season compared to the non-breeding season while males were equally aggressive in both seasons (see Table 1, and Figure 2). There was also a significant interaction effect between sex and treatment (*χ2* = 11.19, *P* = 0.004). While female response to the intruder treatments was overall higher during the non-breeding season than breeding season, they did not differ across intruder treatments in either season (all *P* > 0.2, see Supplementary Table 3). By contrast, males responded significantly more to solo male and duet intruders compared to solo female intruders in both seasons (breeding: *P <* 0.001, *P* = 0.002 respectively; non-breeding: *P =* 0.001, *P* = 0.02). Males responded at similar levels to solo male and duet treatments (breeding: *P =* 0.99; non-breeding: *P =* 0.92).

**Figure 2.**
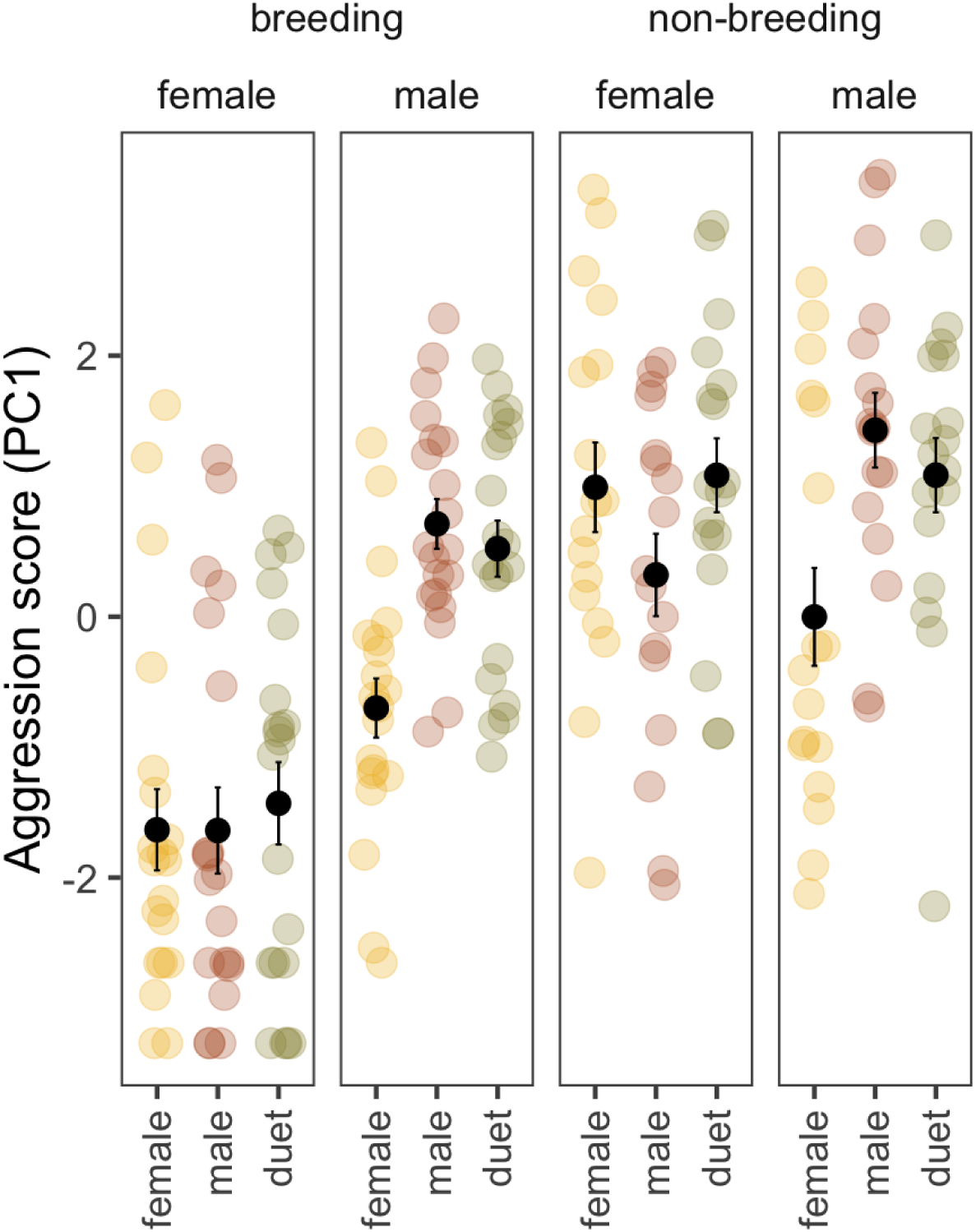
Aggression scores in resident females and males in response to female playback, male playback, and duet playback in the breeding and non-breeding seasons.

**Table 1.** LMM for the aggression score. Bold values indicate statistical significance at alpha set to 0.05.

| *Predictors* | *Estimate* | *95 % CI* | $\chi^2$ | *df* | *p* |
| --- | --- | --- | --- | --- | --- |
| (Intercept) | -1.58 | -2.12 – -1.04 | **33.785** | **1** | **<0.001** |
| treatment [male] | -0.01 | -0.61 – 0.60 | 0.589 | 2 | 0.745 |
| treatment [duet] | 0.20 | -0.41 – 0.81 |  |  |  |
| sex [male] | 0.81 | 0.08 – 1.53 | **4.852** | **1** | **0.028** |
| season [non-breeding] | 2.47 | 1.77 – 3.16 | **49.084** | **1** | **<0.001** |
| treatment [male] × sex [male] | 1.42 | 0.55 – 2.28 | **11.186** | **2** | **0.004** |
| treatment [duet] × sex [male] | 1.02 | 0.16 – 1.88 |  |  |  |
| treatment [male] × season [non-breeding] | -0.67 | -1.57 – 0.23 | 2.460 | 2 | 0.292 |
| treatment [duet] × season [non-breeding] | -0.11 | -1.01 – 0.79 |  |  |  |
| sex [male] × season [non-breeding] | -1.52 | -2.49 – -0.55 | **9.533** | **1** | **0.002** |
| (treatment [male] × sex [male]) × season [non-breeding] | 0.69 | -0.58 – 1.96 | 1.567 | 2 | 0.457 |
| (treatment [duet] × sex [male]) × season [non-breeding] | -0.02 | -1.29 – 1.25 |  |  |  |

#### Singing behavior

All males (N = 20 for the breeding season and N = 17 for the non-breeding season) sang at least one song in response to playbacks during both seasons (see Figure 2a). In contrast, the number of singing females depended on the season (Chi-square test: *χ2*_1_ = 16.91, P < 0.001). Only 15% (3 out of 20) of females responded with song during the breeding season, while 88% (15 out of 17) responded with song during the non-breeding season.

During the non-breeding season, female birds sang most of their songs as duets (proportion of duets to all songs, mean = 0.66, SD = 0.36), whereas males mostly sang solo songs (proportion of duets to all songs, mean = 0.20, SD = 0.24). Of the 169 duets we recorded across trials, 143 (85%) were initiated by males. The number of solo songs produced in response to playback depended on sex, whereby males produced more solo songs than females (*χ2* = 24.97, *P* < 0.001, see Table 2). Females produced more solo songs in response to male intruders than duetting (contrast estimate = 0.65, *P* = 0.006) or female intruders (contrast estimate = 0.96, *P* = 0.0001). Males, on the other hand, sang an equal number of solo songs in response to male and duetting intruders than female intruders (contrast estimates = 0.80 and 0.65, respectively, for male vs. female and duet vs. female comparisons, both *P* < 0.001, see Supplementary Table 4). Although the interaction effect between treatment and sex was statistically not significant (*χ2* = 4.71, *P* = 0.09). For the number of duets, there was an order effect (*χ2* = 7, *P* = 0.03), pairs produced more duets in their third trial compared to the second one (contrast estimate = 0.56, *P* = 0.02, all other comparisons, *n.s.*). Controlling for the influence of trial order, there was a significant effect of treatment (*χ2* = 19.34, *P* < 0.001, see Table 3). Pairs formed more duets in response to male and duet playbacks than female playback (contrast estimates = 0.84 and 1.01, respectively, for male vs. female and duet vs. female comparisons, both *P* < 0.001, see Supplementary Table 5). Duet output of the resident pair to male and duet playbacks were similar (contrast estimate = 0.16, *P* = 0.57)

**Table 2.**
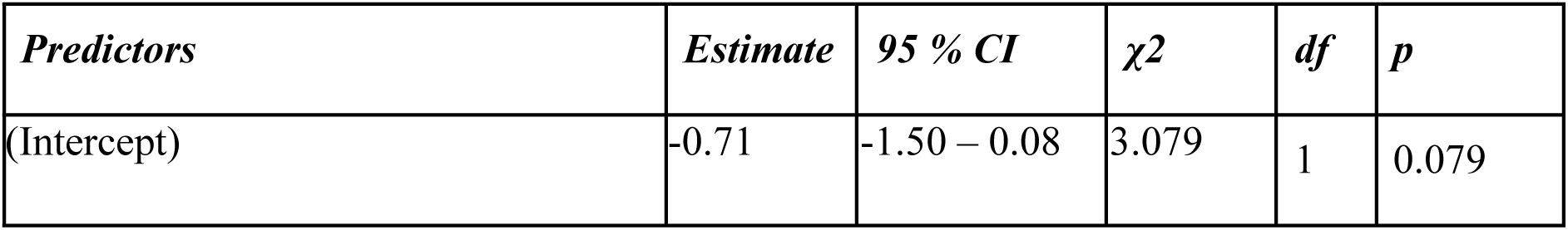
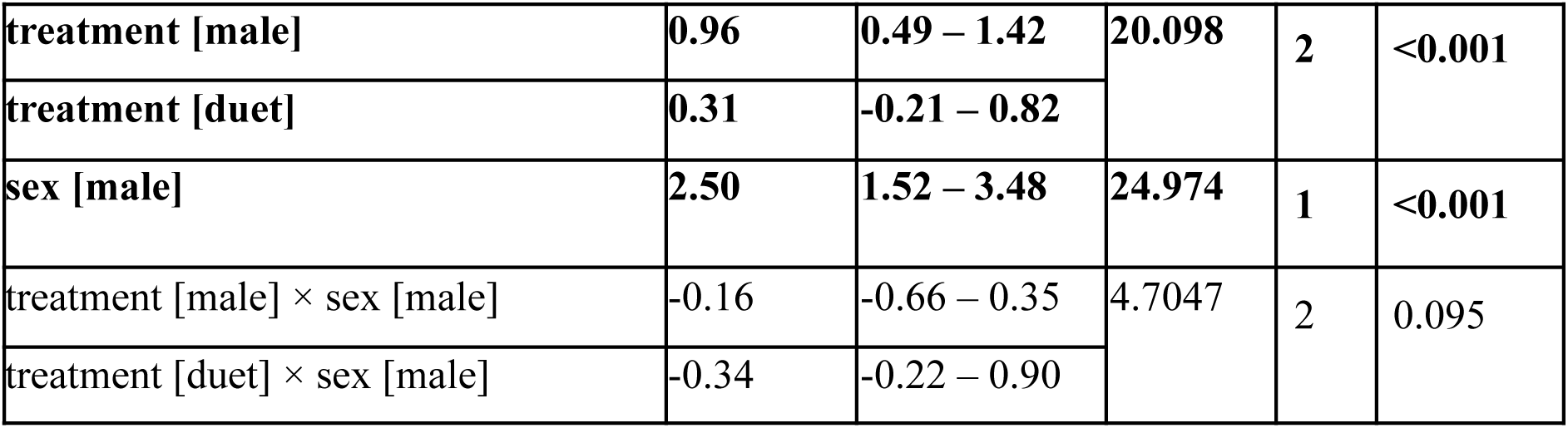
GLMM output for the number of solo songs model. Bold values indicate statistical significance at alpha set to 0.05.

**Table 3.** GLMM output for the number of duets model. Bold values indicate statistical significance at alpha set to 0.05.

| *Predictors* | *Estimate* | *95 % CI* | *χ<sup>2</sup>* | *df* | *p* |
| --- | --- | --- | --- | --- | --- |
| (Intercept) | 0.07 | -0.60 – 0.73 | 0.041 | 1 | 0.839 |
| treatment [male] | **0.84** | **0.38 – 1.30** | **19.342** | **2** | **<0.001** |
| treatment [duet] | **1.01** | **0.56 – 1.46** |  |  |  |
| treatment order [second] | -0.36 | -0.80 – 0.07 | **7.00** | **2** | **0.03** |
| treatment order [third] | **0.20** | **-0.15 – 0.54** |  |  |  |

#### Association between aggression and song in males and females

The association between aggressive behavior during the trials and songs produced after the trial depended on sex (*χ2* = 7.71, *P* = 0.006). In males, song output was positively associated with aggression, whereas in females, there was no such relationship (see Figure 4).

**Figure 3.**
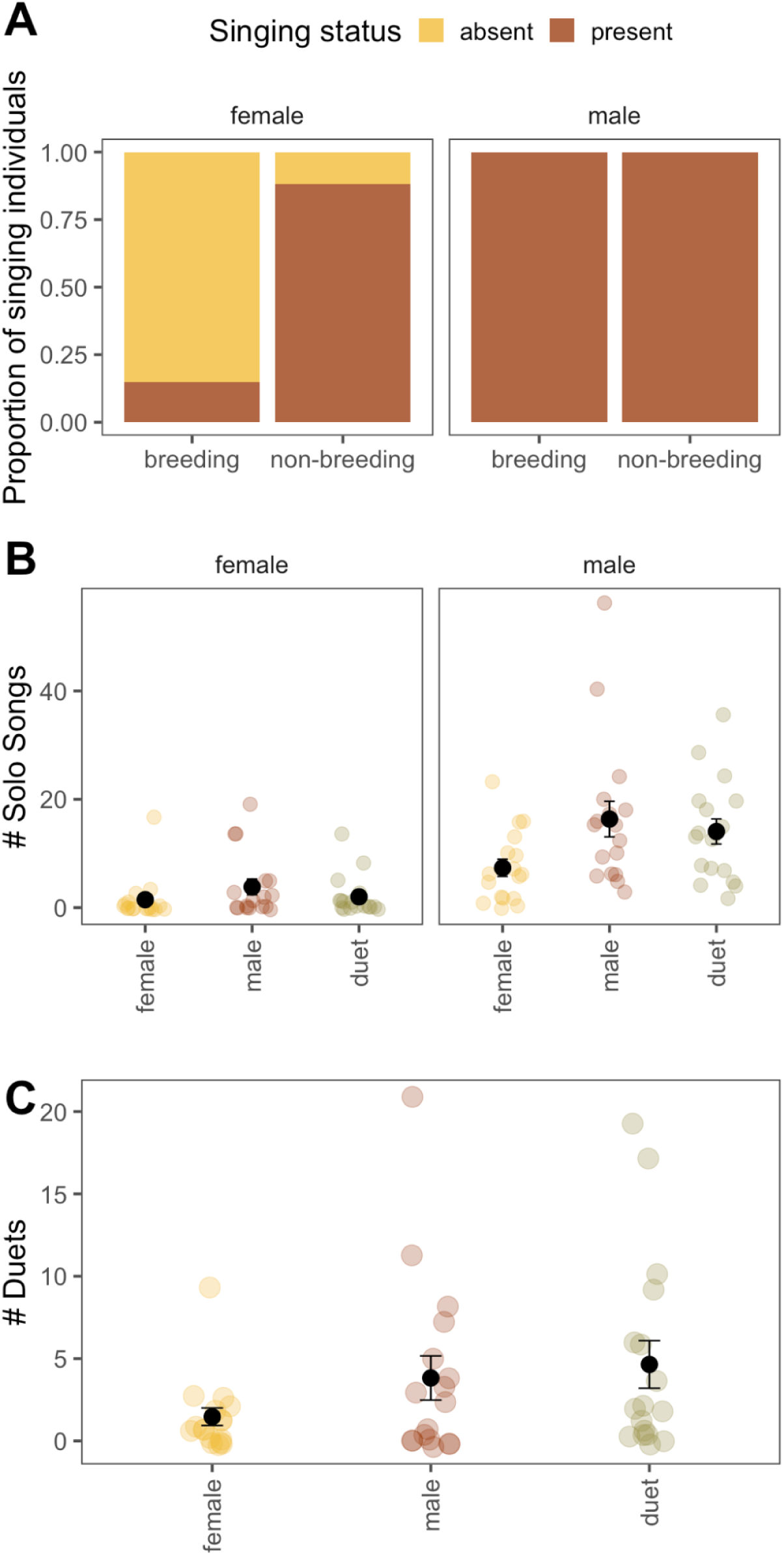
Songs produced in response to playbacks. A) proportion of female and male individuals that sang in response to at least one of the treatments. B) Number of solo songs produced in response to playback treatments by each sex in the non-breeding season. C) Number of duets produced in response to playback treatments by each pair in the non-breeding season.

**Figure 4.**
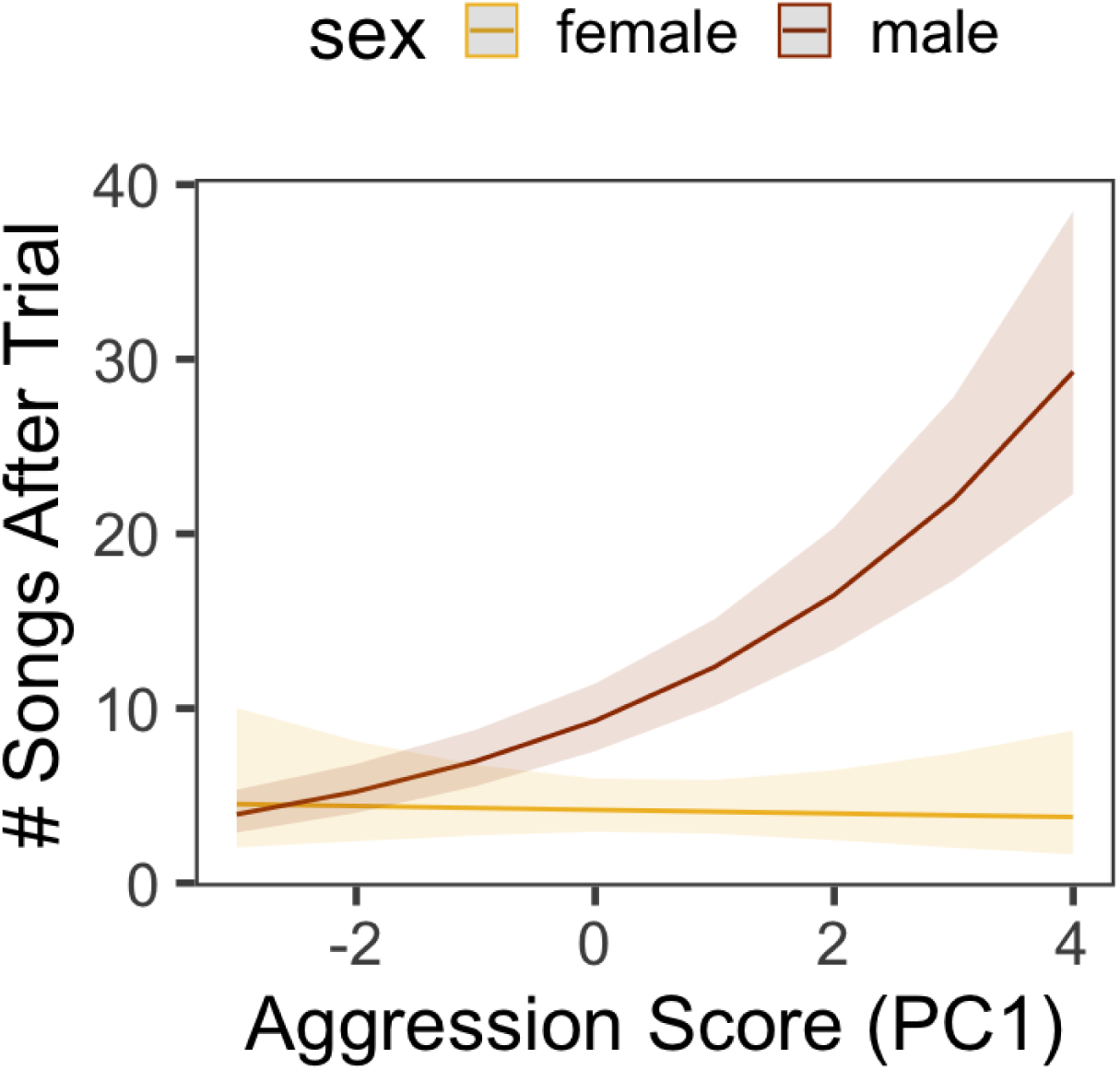
Relationship between the aggression scores (PC1) and song output in the first 5 minutes after the playback, for each sex. Males are represented with red and females are represented with the yellow line. For males, there was a positive relationship between aggression score and song output, however, such a relationship was absent for the females.

### Female song, aggression, and territory retention

Of the 26 color-banded females observed across years, 7 lost their territory (in 2/7 the female was missing while the male was still present, and in 5/7 the pair was missing). For the nine colour-banded females that were tested in the playback experiment, neither the relationship between average song rate and the probability of territory retention (*χ2* = 0.34, *P* = 0.56), nor the relationship between average aggression score and the probability of territory retention was significant (*χ2* = 1.45, *P* = 0.23). Nevertheless, it should be noted that the sample size for these analyses was quite small.

## Discussion

In this study, we tested the function of female song in the Galápagos yellow warbler with a multi-seasonal playback experiment. Our results showed a strong seasonal pattern in female singing and aggression in response to playbacks. Females responded to playbacks by singing and being aggressive, mostly in the non-breeding season. In contrast to the North American subspecies (Hobson & Sealy, 1990), females of the Galápagos subspecies mainly sang duet songs replying to their male partner. Moreover, females sang fewer solo or duet songs towards female song playback than male song or duet playback, providing no support for the intrasexual competition hypothesis for singing. The absence of association between female aggressive behavior and song output in response to playbacks suggests that female song (either solo or duet) does not signal threat in territory defense. Overall, territory loss occurred in seven out of twenty-six females (27%), which is comparable to the proportion observed in females for whom we have systematic playback data, where three out of nine (33%) lost their territories across years. Territory retention in the latter nine females was not explained by song rate or aggression scores in response to playbacks, although we fully acknowledge the small sample size. These findings provide no support for the territorial defense function of female song. In the following paragraphs, we discuss these results in the context of the species’ natural history and other possible functions of female song.

Although both sexes stay together in their territory year-round, and in some cases for multiple years, only the male yellow warblers sang and defended in response to playbacks in both seasons. Females, on the other hand, mainly sang and defended in the non-breeding season. This discrepancy between sexes might be due to differential costs of singing. The same pattern was observed in the sedentary rufous-capped warbler (*Basileuterus rufifrons*); females sang less often and were less aggressive in response to intruders during the breeding season, while males showed strong responses in both seasons (Demko & Mennill, 2018). In both species, females are the primary contributors of nest building and incubation, which may pose opportunity costs to singing and participating in territory defense. Alternatively, females might be keeping quiet due to higher relative nest predation risks of singing during breeding. For instance, Kleindorfer and colleagues (2016) found that in superb fairy-wrens (*Malurus cyaneus*), female song rate predicted offspring predation rates likely because females were singing closer to (and sometimes inside) the nest than males.

Hobson and Sealy (1990) suggested that female song in the North American yellow warbler mainly functions in intrasexual competition for resources because they observed female song prior to nest building; also, 2 out of 4 females that they recorded sang in response to a female decoy in their territory. Under this hypothesis, we predicted that females would be more aggressive and sing more towards female intruders. However, our results did not support this. During the non-breeding season (when the female song was mainly present), females were equally aggressive towards all intruders, but they sang the least amount of duet and solo songs towards female intruders. This difference is paralleled in the singing patterns of the two subspecies: females only were observed to sing solo songs in North America but females sang mostly duets in Galápagos. In a meta-analysis, Logue (2005) found that same-sex aggression bias was more common in non-duetting species, while duetting ones were more cooperative in defense, showing less bias towards the same sex. Nevertheless, experimental support for the intrasexual competition hypothesis in the North American species is also lacking.

While aggression predicted song output in males, female aggression did not predict song output in response to intruder playbacks. This finding provides no support for the territory defense function of female song. Although territory defense may be the most common function of female song in the systems studied so far, it is not the case for all species (Barros et al., 2024). Eastern bluebird females didn’t increase song rates in response to intruders, even though they were being aggressive toward intruders; instead, female songs were mainly used as an intrapair communication signal during intrusions (Rose et al., 2019). Similarly, our study design doesn’t allow us to rule out the possibility that the female song might be used in the presence of intruders as a pair-directed signal to coordinate defense. Female yellow warblers could be singing to signal their location to their pair during an intrusion, either to increase proximity or prevent misdirected aggression (Dahlin & Benedict, 2014).

We had predicted that duetting pair intruders would receive the strongest aggressive response from both sexes; however, this was not the case. Females were equally aggressive towards all intruders, and males were equally and more aggressive towards males and duetting intruders than the female intruders. It is possible that the use of a single speaker for a duetting pair was perceived as unnatural (Koloff & Mennill, 2013). Nonetheless, given that we often observed pairs duetting as close as <1m to each other in our study population, we don’t believe the experimental set-up was unrealistic. Instead, we speculate that if a solo male intruder already poses a very high threat, the pair might already be showing aggression at their maximum capacity, resulting in a ceiling effect.

Given the lack of support for either the intrasexual competition or the territory defense hypothesis, we believe that the most likely hypothesis for the function of female song and duetting in Galápagos yellow warblers is that it is a mate-directed signal, used to coordinate the social activity of the pair. More specifically, pairs may form duets to maintain contact with each other (Hall, 2004). In this case, we would expect pairs to be visually separated when duetting. Additionally, as noted in the previous paragraphs, duets may facilitate cooperation during territory defense. Under this hypothesis, we expect pair proximity during defense to correlate with duet output. We are currently testing these hypotheses experimentally as well as with observational data.

In summary, we show that female song is very common in a year-round resident subspecies of a widely studied species, the yellow warbler, 35 years after the first report by Hobson and Sealy (1990). This adds to recent studies of widely studied species where female song is only now described to be common (Sierro et al., 2022, blue tit, *Parus caeruleus*; Wilkins et al., 2020, barn swallow, *Hirundo rustica*). Our results reveal the importance of studying bird song in multiple seasons and of evaluating the function of female song as well as male song, and considering multiple hypotheses.

## Data availability

Data and R script to reproduce analyses can be found in the supplementary materials.

## Acknowledgments

Permission to conduct this study was granted by the Galápagos National Park Directorate (permit numbers PC-87-23 and PC-05-25) with logistical support provided by the Charles Darwin Research Station (CDRS). We are grateful to the Floreana community for their support. We thank Jefferson García Loor, Lauren Common, Andrew Katsis, Melanie Kaluppa, Leon Hohl, Alena Hohl, Matt Millham, Caelan Linke, Abbie Hay, Johanna Kniely, Tim Garret, Rachel Dudaniec, Ursula Scuderi, and Roland Digby for their help with fieldwork. We thank Lena Gies for helpful feedback on the manuscript and providing the illustrations. This publication is contribution number -to be filled upon acceptance- of the Charles Darwin Foundation for the Galápagos Islands. This work was funded by the Austrian Science Fund (Project numbers W1262-B29 [10.55776] and P 36342-B) with awards to SK.

## Author contributions

AY, CA, and SK designed the research; AY, CA and KAM conducted the playback trials; AY analysed the data and wrote the first draft of the manuscript; all authors revised the manuscript and gave the final approval for publication.

