## Supplementary Materials for "Solo songs, duets, and territory defence across seasons in female Galápagos Yellow Warblers (*Setophaga petechia aureola*)"

**Supplementary Table 1.** Number of pairs that received treatments in the corresponding order in each season

| **Treatment Order** | **Non-breeding Season** | **Breeding Season** |
| --- | --- | --- |
| Male-Female-Duet | 3 | 3 |
| Male-Female-Duet | 3 | 3 |
| Female-Male-Duet | 3 | 4 |
| Female-Male-Duet | 3 | 3 |
| Male-Duet-Female | 3 | 3 |
| Male-Duet-Female | 2 | 3 |

**Supplementary Table 2.** Factor loadings to PC1 (Aggression Score) for each variable.

| **Response variable** | **Factor loading** |
| --- | --- |
| **Flights** | -0.51 |
| **Time spent within 5m of the speaker** | -0.50 |
| **Closest approach to the speaker** | 0.53 |
| **Latency to respond to playback** | 0.44 |

**Supplementary Table 3.** Post-hoc pairwise comparisons of the aggression score LMM. Bold values indicate statistical significance at alpha set to 0.05.

| **Contrast** | **Season** | **Estimate** | **SE** | **df** | **t** | ***p*** |
| --- | --- | --- | --- | --- | --- | --- |
| female female - male female | breeding | 0.005 | 0.309 | 154.677 | 0.017 | 1 |
| female female - duet female | breeding | -0.203 | 0.309 | 154.677 | -0.656 | 0.986 |
| female female - female male | breeding | -0.809 | 0.368 | 115.864 | -2.197 | 0.247 |
| **female female - male male** | **breeding** | **-2.22** | **0.368** | **115.864** | **-6.026** | **0** |
| **female female - duet male** | **breeding** | **-2.031** | **0.368** | **115.864** | **-5.512** | **0** |
| male female - duet female | breeding | -0.208 | 0.309 | 154.677 | -0.673 | 0.985 |
| male female - female male | breeding | -0.814 | 0.368 | 115.864 | -2.21 | 0.241 |
| **male female - male male** | **breeding** | **-2.225** | **0.368** | **115.864** | **-6.04** | **0** |
| **male female - duet male** | **breeding** | **-2.036** | **0.368** | **115.864** | **-5.526** | **0** |
| duet female - female male | breeding | -0.606 | 0.368 | 115.864 | -1.646 | 0.57 |
| **duet female - male male** | **breeding** | **-2.017** | **0.368** | **115.864** | **-5.476** | **0** |
| **duet female - duet male** | **breeding** | **-1.828** | **0.368** | **115.864** | **-4.961** | **0** |
| **female male - male male** | **breeding** | **-1.411** | **0.309** | **154.677** | **-4.569** | **0** |
| **female male - duet male** | **breeding** | **-1.221** | **0.309** | **154.677** | **-3.955** | **0.002** |
| male male - duet male | breeding | 0.19 | 0.309 | 154.677 | 0.614 | 0.99 |
| female female - male female | non-breeding | 0.672 | 0.335 | 154.677 | 2.006 | 0.344 |
| female female - duet female | non-breeding | -0.092 | 0.335 | 154.677 | -0.273 | 1 |
| female female - female male | non-breeding | 0.711 | 0.397 | 126.003 | 1.792 | 0.475 |
| female female - male male | non-breeding | -0.719 | 0.397 | 126.003 | -1.813 | 0.461 |
| female female - duet male | non-breeding | -0.375 | 0.397 | 126.003 | -0.946 | 0.934 |
| male female - duet female | non-breeding | -0.763 | 0.335 | 154.677 | -2.279 | 0.209 |
| male female - female male | non-breeding | 0.039 | 0.397 | 126.003 | 0.098 | 1 |
| **male female - male male** | **non-breeding** | **-1.391** | **0.397** | **126.003** | **-3.507** | **0.008** |
| male female - duet male | non-breeding | -1.047 | 0.397 | 126.003 | -2.639 | 0.095 |
| duet female - female male | non-breeding | 0.802 | 0.397 | 126.003 | 2.022 | 0.336 |
| duet female - male male | non-breeding | -0.628 | 0.397 | 126.003 | -1.583 | 0.612 |
| duet female - duet male | non-breeding | -0.284 | 0.397 | 126.003 | -0.715 | 0.98 |
| **female male - male male** | **non-breeding** | **-1.43** | **0.335** | **154.677** | **-4.27** | **0** |
| **female male - duet male** | **non-breeding** | **-1.086** | **0.335** | **154.677** | **-3.242** | **0.018** |
| male male - duet male | non-breeding | 0.344 | 0.335 | 154.677 | 1.028 | 0.908 |

**Supplementary Table 4.** Post-hoc pairwise comparisons of the solo song model. Bold values indicate statistical significance at alpha set to 0.05.

| **Contrast** | **Sex** | **Estimate** | **SE** | **z ratio** | ***p*** |
| --- | --- | --- | --- | --- | --- |
| female - male | female | -0.956 | 0.235 | -4.060 | 0.0001 |
| female - duet | female | -0.307 | 0.263 | -1.167 | 0.4729 |
| male - duet | female | 0.648 | 0.212 | 3.062 | 0.0062 |
| female - male | male | -0.799 | 0.108 | -7.422 | <.0001 |
| female - duet | male | -0.648 | 0.110 | -5.872 | <.0001 |
| male - duet | male | 0.151 | 0.088 | 1.714 | 0.2000 |

**Supplementary Table 5.** Post-hoc pairwise comparisons of the duet model. Bold values indicate statistical significance at alpha set to 0.05.

| **Contrast** | **Estimate** | **SE** | **z ratio** | ***p*** |
| --- | --- | --- | --- | --- |
| **female - male** | **-0.841** | **0.236** | **-3.568** | **<0.001** |
| **female - duet** | **-1.010** | **0.231** | **-4.372** | **<0.001** |
| male - duet | -0.170 | 0.168 | -1.012 | 0.569 |

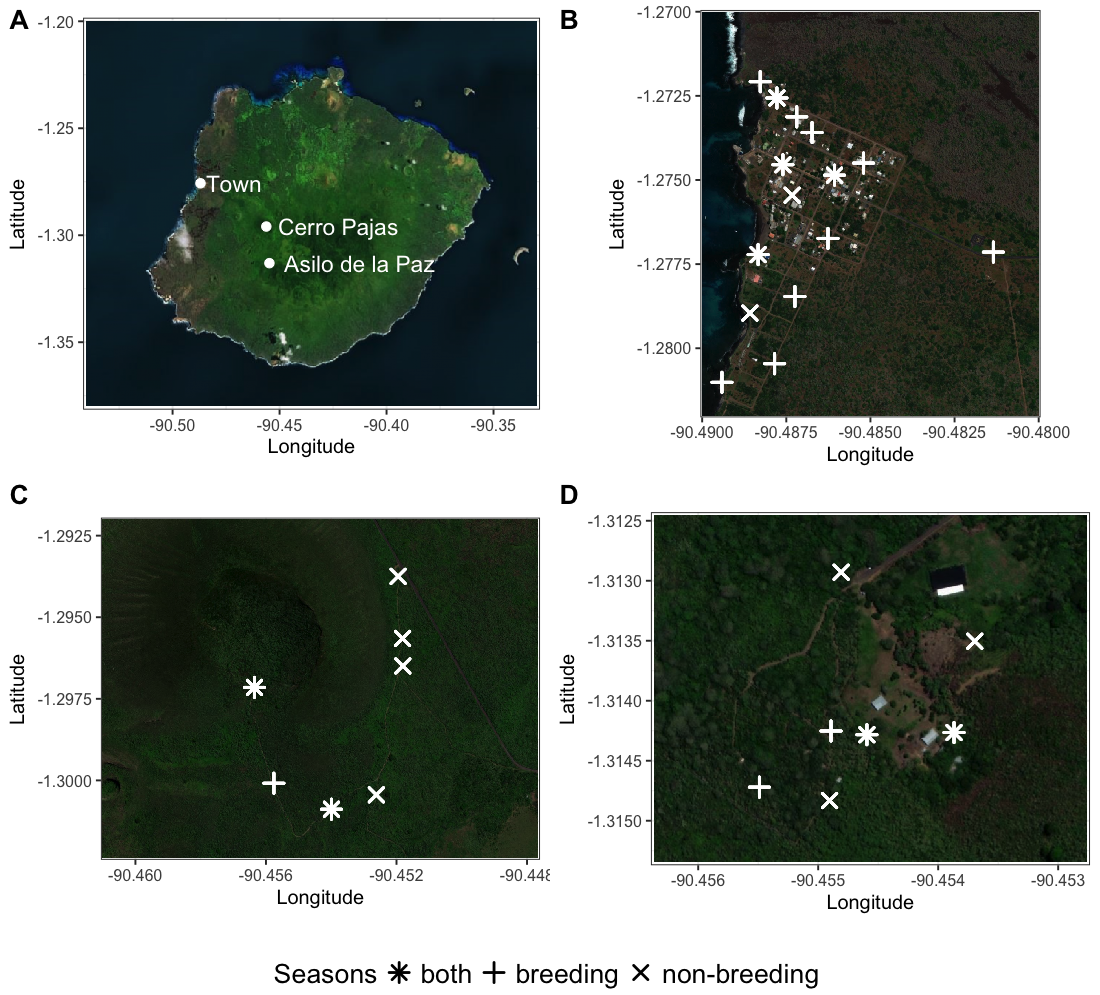

**Supplementary Figure 1.** Satellite maps of study sites and locations of the playback experiments. A) Three study sites are shown on Floreana Island B) Town C) Cerro Pajas D) Asilo de la Paz. Each study territory is shown on maps B, C, and D with white markers, their shape indicates the season in which the experiments were done.
